# Exploring the relationship between altmetrics and traditional measures of dissemination in health professions education

**DOI:** 10.1101/260059

**Authors:** Lauren A. Maggio, Todd C. Leroux, Holly S. Meyer, Anthony R. Artino

**Author notes:** Corresponding author Lauren Maggio 4301 Jones Bridge Road, Bethesda, MD 20814 USA. Twitter: @laurenmaggio. Disclaimer: The views expressed in this article are those of the authors and do not necessarily reflect the official policy or position of the Uniformed Services University of the Health Sciences, the U.S. Department of Defense, or the U.S. Government.

## Abstract

Researchers, funders, and institutions are interested in understanding and quantifying research dissemination and impact, particularly related to communicating with the public. Traditionally, citations have been a primary impact measure; however, citations can be slow to accrue and focus on academic use. Recently altmetrics, which track alternate dissemination forms (e.g., social media) have been suggested as a complement to citation-based metrics. This study examines the relationship between altmetrics and traditional measures: journal article citations and access counts.

The researchers queried Web of Science and Altmetric Explorer for articles published in HPE journals between 2013-2015. They identified 2,486 articles with altmetrics. Data were analyzed using negative binomial and linear regression models.

Blogging was associated with the greatest increase in citations (13% increase), whereas Tweets (1.2%) and Mendeley (1%) were associated with smaller increases. Journal impact factor (JIF) was associated with a 21% increase in citations. Publicly accessible articles were associated with a 19% decrease, but the interactive effect between accessible articles and JIF was associated with a 12% increase. When examining access counts, publicly accessible articles had an increase of 170 access counts whereas blogging was associated with a decrease of 87 accesses.

This study suggests that several altmetrics outlets are positively associated with citations, and that public accessibility, holding all other independent variables constant, is positively related to article access. Given the scientific community’s evolving focus on dissemination—including to the public—these findings have implications for stakeholders, providing insight into the factors that may improve citations and access of articles.

## Introduction

Research dissemination is essential for scientific progress. As part of their commitment to scholarly work, researchers have a duty to disseminate their research as widely as possible to all those who are interested in and might benefit from it[1]. This responsibility is even more apparent when one considers the large proportion of scientific research that is funded with public money.

Underscoring this commitment, in 2008 the National Institutes of Health (NIH), the largest public funder of biomedical research in the world[2], adopted the NIH Public Access Policy[3]. This policy mandates that grant recipients disseminate and make publicly accessible any peer-reviewed publications resulting from NIH-funded research within 12 months of publication. If investigators fail to comply, their research funds may be frozen and future funding applications jeopardized.

More recently, the NIH expanded its dissemination efforts to encourage researchers to publicly deposit data resulting from such work to ensure access by the public, the scientific community, and industry[4]. Globally, in 2017, more than 80 major funders (e.g., Canadian Institutes of Health Research, European Research Council, Brazilian Ministry of Science and Technology) followed suit requiring grantees to make funded research publicly accessible[5]. The rationale for these policies is predicated on a fundamental principle: progress in science requires access to and dissemination of research results.

Traditionally, research dissemination – and ultimately, research uptake – have been measured by article-and journal-level metrics such as citation counts and journal impact factor (JIF), a citation-based metric. Referred to as “pellets of peer recognition”[6], citations have been criticized for accumulating slowly and not reflecting societal attention from outside of academia[7-9]. On the other hand, JIF has been disparaged for being opaque in its methods, skewed by the relatively few articles in a journal that receive the bulk of citations, and easily gamed[10, 11]. Furthermore, JIF has been charged with causing many scientists to focus too much on publishing in “high-impact journals” and not enough on doing high-quality science with societal impact[12]. Finally, a citation-focused approach does not address the need to reach the public[4], and there is some indication that traditional metrics are poor measures of practical impact in applied fields such as clinical medicine[9].

In light of these limitations, the scientific community has become interested in other ways to capture research dissemination and impact, particularly as it relates to communicating with the general public. Moreover, researchers, publishers, funders, and academic institutions have started to take seriously the benefits of sharing their research outside of academia, using alternative channels, such as the news media and social media platforms like Twitter, Facebook, and Instagram. As this dissemination approach has become more popular, the scientific community has turned to alternative metrics, or altmetrics, to measure the degree to which their research is shared and discussed on non-traditional channels-something that is not captured by traditional citation-based metrics[13, 14].

Altmetrics have been described as “web-based metrics for the impact of scholarly material, with an emphasis on social media outlets as sources of data”[15]. In other words, altmetrics “provide a summary of how research is shared and discussed online, including by the public”[16]. In this way, altmetrics are seen as complementing, not replacing, traditional citation-based metrics[17].

While some scholars believe altmetrics are a valuable way to measure dissemination and scholarly impact[16, 18], others question if they may be “just empty buzz" [15]. This concern stems, in part, from the idea that researchers can potentially “game the system” (e.g., by tweeting or blogging about their own work to bolster their article’s online attention). In addition, other scholars argue that a research paper’s social media popularity does not distinguish between positive and negative attention and therefore may be a poor proxy for quality and unrelated to other dissemination metrics. On this point, prior research exploring the links between altmetrics and traditional metrics is mixed. A recent *PLOS ONE* study of ecology research found that after controlling for JIF, Twitter activity was one of the best predictors of citation rates, second only to time since publication[19]. On the other hand, two larger studies conducted across multiple disciplines (including social science disciplines) found only weak, though still positive, correlations between social media activity and traditional citation rates[17, 20]. Results from these latter studies suggest that altmetrics may be assessing something different than citation rates.

## Altmetrics of health professions education

Health professions education (HPE) is a field that “aims to supply society with a knowledgeable, skilled, up-to-date cadre of professionals who put patient care above self-interest and undertake to maintain and develop their expertise over the course of a lifelong career” [21]. In HPE, professionals encompass a range of practitioners including, but not limited to, physicians, nurses, dentists, and pharmacists. As a relatively new and highly applied field, there is reason to believe the links between altmetrics and more traditional metrics might differ for HPE research articles, as compared to other fields[9, 22].

There is limited research on altmetrics in HPE, despite demonstrably strong growth in the number of altmetrics events for articles appearing in HPE-focused journals[16] and growing interest from HPE researchers[18]. Currently, there is only one study of altmetrics conducted with HPE articles published in the journal *Medical Education*; it found weak to moderate correlations between altmetrics, access counts, and citations for articles published in 2012 and 2013[22]. Unfortunately, altmetrics studies of biomedical research have generally not included HPE research[23]. Considering the limited scope of prior work in HPE research, as well as the speed at which social media technology is changing the research dissemination landscape [16, 24], more generalizable work is needed to better understand how altmetrics relate to traditional article-level metrics across HPE research.

Therefore, the purpose of this article is to examine the relationship between altmetrics and two traditional measures of dissemination and impact in HPE: journal article citations and article access counts. Exploring these relationships is important for at least two reasons. First, this work can inform the community by modernizing our understanding of scholarly dissemination and impact in HPE. And second, from a practical standpoint, it can help individual HPE scholars determine the best ways to use social media and other non-traditional outlets to disseminate and promote their research to academic colleagues and the general public.

## Methods

This cross-sectional, quantitative, bibliometric study was designed to examine the relationship between altmetrics events, article citations, and article access counts. To address our primary aim, we used aggregated data with no personal identifiers. As such, this study did not require ethics approval.

## Data collection

We searched Web of Science (WOS) on 2/22/2017 for articles published in the following seven HPE journals: *Academic Medicine, Medical Education, BMC Medical Education, Advances in Health Sciences Education, Medical Teacher, Teaching and Learning in Medicine and Journal of Continuing Education in the Health Professions*. We based our selection on previous research that had identified these journals as key publications in HPE[25, 26]. Additionally, we limited the sample to journals with an impact factor for all years studied as we wanted to control for JIF in our models. The search was restricted to articles published from 2013 to 2015 and to research articles as defined by WOS. All retrieved article citations, including available metadata, were downloaded into an Excel spreadsheet. Metadata included the number of times an article was cited.

To determine if articles retrieved from WOS had an altmetrics event (a tweet, media mention, Mendeley save, etc.), we searched Altmetric Explorer (Altmetric, London UK) on May 17, 2017 for articles with altmetrics events published in the specified HPE journals. Altmetric Explorer is a search tool that queries a database of over 7 million articles that have had an altmetrics event in at least one of sixteen altmetrics outlets sourced by Altmetric[27]. For further information on Altmetric’s data collection strategies, please see: https://www.altmetric.com/about-our-data/how-it-works/. We merged retrieved article references from Altmetric Explorer with WOS data in the existing Excel file. Articles identified by the Altmetric Explorer search, but not retrieved by WOS, were discarded (e.g., commentaries, letters to the editor). Thus, our overall data set was limited to research articles. We retained articles in the WOS data set that did not include altmetrics events, and entered zeroes for such articles for all altmetrics fields.

## Access data

We obtained article access counts for BMC *Medical Education, Advances in Health Sciences Education, Medical Teacher, and Teaching and Learning in Medicine* by visiting each article’s abstract page on the publisher’s website and noting the number of times the article’s full-text was accessed. Access counts include full-text article accesses by users that directly visited the journal’s website or who arrived there via a database or search engine (e.g., PubMed, library database, or Google). Data were extracted over three days (4/9/2017-4/11/2017) and added to the master Excel spreadsheet.

Access count information was not publicly available for *Academic Medicine, Journal of Continuing Education in Health Professions*, and *Medical Education*. Therefore, we requested the data from each journal’s editor. Access counts from *Academic Medicine* and *Medical Education* were provided on 5/18/2017 and 5/19/2017, respectively, and included HTML and PDF views. We were unable to obtain access counts from the *Journal of Continuing Education in the Health Professions*, and thus did not include its articles in our study. This journal had 109 article citations with an altmetrics event, which accounted for approximately 4 percent of the overall sample.

When collecting access counts, we noted for each article if it was publicly accessible. We defined an article to be publicly accessible if the full-text of the article was freely available to read without the need for a subscription or payment to the publisher’s website. Thus, publicly accessible articles included those that have open access (OA) licenses (e.g., CC BY, CC BY NC) and “bronze" articles, which were free to read on a publisher’s website but did not include an explicit OA license[28]. For the three journals without readily available citation counts, we accessed each citation on its respective journal’s website to determine its public accessibility status. Of note, for our study, all *Academic Medicine* articles were bronze OA, in that the journal makes all articles that are one year post-publication freely available on its website. Additionally, *BMC Medical Education* is an OA journal, and therefore all of its articles include an OA license and are publicly accessible from the time they are published.

## Statistical analysis

In addition to descriptive and summary statistics, two regression models were used to evalúate the relationship between altmetrics outlets and citations and access count statistics. The first model consisted of a negative binomial regression with citations as the dependent variable, and the second model was an ordinary least squares linear regression model with access counts as the dependent variable. Separate models were employed due to the underlying distributions of the dependent variables (i.e., the dependent variable in the first model was not normally distributed) Adequate model fit was assessed using criterion fit statistics and examination of residual errors for any unusual, systematic behavior. The independent variables for the models included time since publication (measured in weeks), JIF for the specific journal at the time the article was published, whether the article was publicly accessible (dummy variable), the number of references in the article, activity related to Twitter, Facebook, news outlets, blog posts, Mendeley saves, and an interaction between JIF and whether the article was publicly accessible. The interaction was included in both models because we suspected that journals of varying JIFs might operate differently when it comes to whether or not the article is publicly accessible. We focused on Twitter, Facebook, news outlets, blogs, and Mendeley saves as these were previously identified as the most prevalent altmetrics outlets in HPE research[16]. The statistical analysis was performed using R[29], with packages that include ggplot2[30], gridExtra[31], psych[32], reshape[33], and COUNT[34]. Data and code for this project are available at https://github.com/toddleroux/health_prof_educ.

## Results

We analyzed 2,486 articles. Table 1 presents descriptive and summary statistics for the study sample. The median number of citations was 3, while the average access statistic was 81.5. Journal articles in the sample were evenly distributed across the years 2013 through 2015. The average JIF was 2.3 and the average number of references in each article was 33.0. Concerning altmetrics events, on average, there were approximately 6 Tweets per article, although there was great variability in this statistic. All examined altmetrics outlets were represented. Almost 60 percent of the articles were publicly accessible. For the study sample, activity related to news outlets and blogs was exceptionally low, but Mendeley saves were considerably higher, averaging 20.8.

**Table 1:**
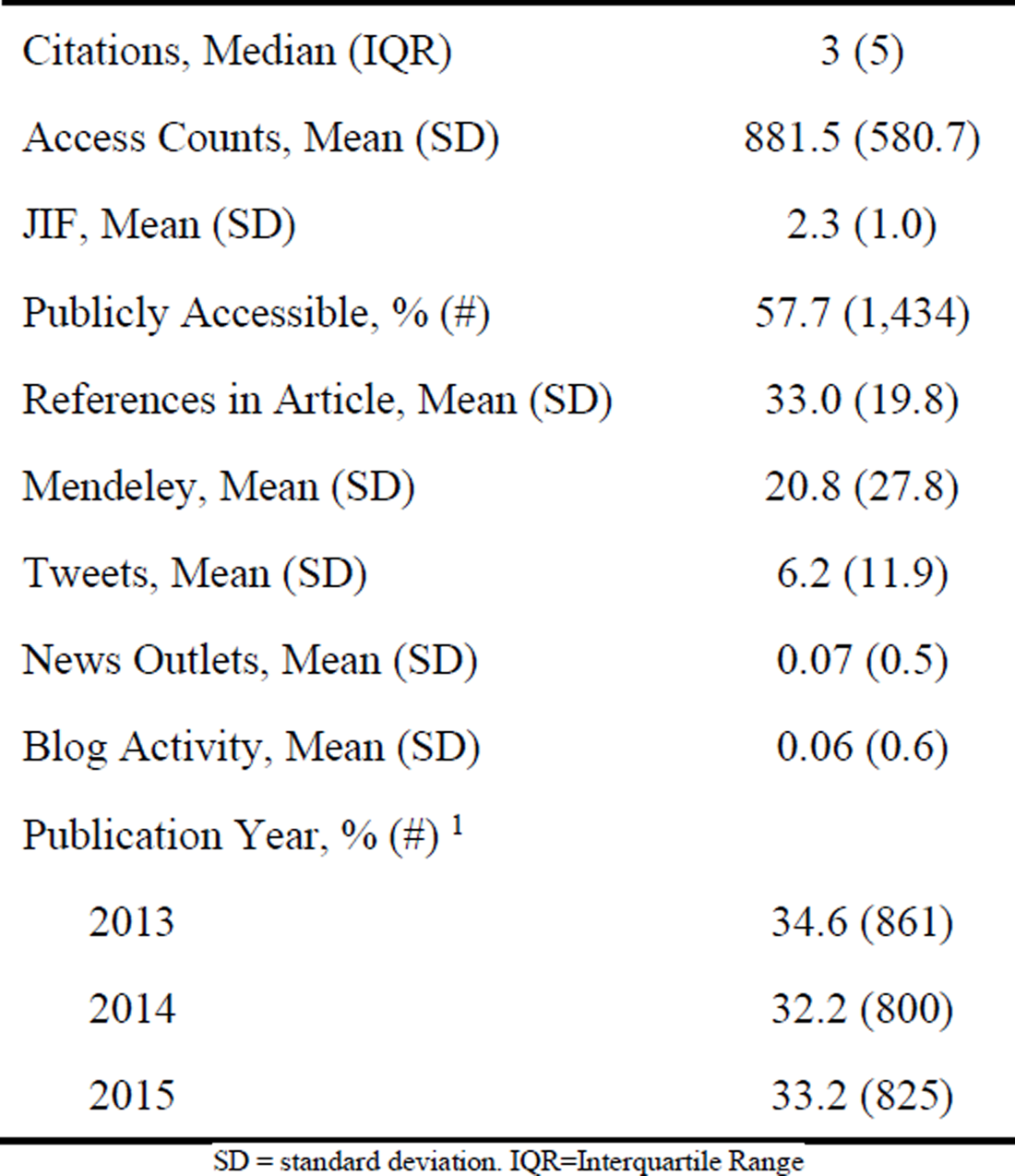
Descriptive and Summary Statistics of Study Sample (n = 2,486)

Table 2 contains model results for the negative binomial regression model whose dependent variable was citations. The coefficient estimates are presented as incident rate ratios, which are the exponent of the original model coefficient estimates. Blogging activity (incident rate ratio [IRR] = 1.13, 95% IRR confidence interval [CI] = 1.01, 1.25) and JIF (IRR = 1.21, 95% IRR CI = 1.13, 1.30) had the largest, positive effect on citations. For example, our model suggests that for each additional blog about an article, citations, on average, would increase approximately 13 percent. Tweets and Mendeley saves were both statistically significantly related to citations, although the effect sizes were small [Tweets (IRR = 1.01, 95% IRR CI = 1.00, 1.01), Mendeley (IRR = 1.01, 95% IRR CI = 1.00, 1.01)]. Publicly accessible manuscripts had a decrease of 19 percent in citation rates (IRR = 0.81, 95% IRR CI = 0.67, 0 The interactive effect of a publicly accessible journal article and JIF was associated with a 12 percent increase in citation rate, on average, as compared to other journals with a non-accessible article (IRR = 1.12, 95% IRR CI = 1.04, 1.21). As the model indicates, the number of references contained in a journal article was statistically significantly associated with, but did not have any practical effect on, an article’s citation rate.

**Table 2:** Citation Negative Binomial Model Results

| Coefficient | IRR | 95% IRR CI |
| --- | --- | --- |
| (Intercept) *** | 0.28 | 0.23, 0.35 |
| Publicly Accessible * | 0.81 | 0.67, 0.98 |
| References *** | 1.00 | 1.00, 1.01 |
| Bloggers * | 1.13 | 1.01, 1.25 |
| Tweets *** | 1.01 | 1.00, 1.01 |
| News Outlets | 0.98 | 0.93, 1.04 |
| Facebook | 0.97 | 0.92, 1.02 |
| Mendeley *** | 1.01 | 1.00, 1.01 |
| Journal Impact Factor *** | 1.21 | 1.13, 1.30 |
| Time (in weeks) *** | 1.01 | 1.00, 1.01 |
| Publicly Accessible × JIF ** | 1.12 | 1.04, 1.21 |

IRR = incident rate ratio, CI = confidence interval, \* = p-value < 0.05, \*\* = p-value < 0.01, \*\*\* = p-value < 0.001.

Table 3 contains linear regression model results for access count statistics. As indicated, a publicly accessible article, on average, had approximately a 169 point increase in access count as compared to a closed journal article (169.62, 95% CI = 49.03, 290.19). Holding all other variables constant, a publicly accessible article would on average garner an extra 169 full-text accesses as compared to an article that is not publicly accessible. Contrary to our first model, the number of references had a small, negative association with usage statistics (−1.52, 95% CI = −2.72, −0.32). Additionally, altmetrics activity on Mendeley savs, Twitter, Facebook, and news outlets did not demonstrate any statistically significant associations with access counts. Blogging activity was negatively associated with access counts (−86.51, 95% CI = −166.12, −6.90). The JIF of the journal was positively related to access counts (143.48, 95% CI = 99.32, 187.63), and time had a very small, positive effect on access counts. As time was measured in weeks, for every one week increase in the time since the article was published, access counts increased, on average, 1.08 (95% CI = 0.56, 1.60). Lastly, and in contrast to the first model, the combination of JIF and public accessibility was negatively associated with access counts, as compared to articles that were not publicly accessible (−89.16, 95% CI = −140.82, −37.51).

**Table 3:** Access Count Linear Regression Model Results

| Coefficient | Estimate | 95% CI |
| --- | --- | --- |
| (Intercept) *** | 455.73 | 322.79, 588.67 |
| Publicly Accessible ** | 169.62 | 49.03, 290.19 |
| References * | -1.52 | -2.72, -0.32 |
| Bloggers * | -86.51 | -166.12, -6.90 |
| Tweets | 0.89 | -1.43, 3.21 |
| News Outlets | -4.10 | -45.11, 36.92 |
| Facebook | -6.63 | -43.40, 30.14 |
| Mendeley | 0.74 | -0.17, 1.67 |
| Journal Impact Factor *** | 143.48 | 99.32, 187.63 |
| Time (in weeks) *** | 1.08 | 0.56, 1.60 |
| Publicly Accessible × JIF *** | -89.16 | -140.82, -37.51 |

\* = p-value < 0.05, \*\* = p-value < 0.01, \*\*\* = p-value < 0.001. Adjusted R-squared: 0.02006 F-statistic: 7.032 on 11 and 3231 DF, p-value: 6.824e-12.

## Discussion

Broad dissemination of research is essential for continued growth of HPE. Moreover, researchers’ knowledge of the dissemination opportunities has the potential to improve the scope and breadth of their work’s reach. Therefore, we analyzed the relationship between altmetrics and journal article citations and article access counts. To our knowledge, this is the first study focused on HPE research that has incorporated more than one journal, spanned across several years, and considered the effects of public accessibility.

## Citations

Despite criticisms, citations are considered a proxy for impact in academia. Thus, opportunities to increase citation rates can be advantageous for researchers, especially those seeking tenure and promotion[18, 35]. In our model, JIF was associated with the largest citation increase. Moreover, Tweets and Mendeley saves were also related to a citation increase (albeit very small increases). This finding aligns with a broader study of PubMed articles that found a citation advantage for Tweets (Mendeley was not included in this previous analysis)[36]. In addition, the public accessibility of an article was related to a fairly large decrease in citation rates (19 percent), a surprising finding that does not align with previous research [35, 40, 41].

For HPE, our findings suggest that researchers and journal editors concerned with citations might consider utilizing altmetrics outlets to communicate findings. In some cases, HPE researchers and journals have already embraced altmetrics outlets, especially social media, with HPE authors tweeting articles (generally with #meded or #hpe) and journals posting article links to their Facebook pages[37].

A citation advantage was notable for articles that were blogged. This finding aligns with previous research[17, 38] and is unsurprising given the fairly strong presence of blogs in medical education[39]. That said, this finding stands contrasts with previous HPE study that did not identify a blog post advantage[22]. It is worth noting, however, that this previous study was conducted on a single journal. Still, these contradictory findings suggest the need for further research on the relationship between blogs and citations, taking into account the characteristics of included journals. For example, researchers might examine differences in journals that have their own dedicated blog (or professional association that maintains a journal blog feature) and those that do not.

In the present study, we did not identify a citation advantage for publicly accessible HPE articles. We were surprised by this finding since we assumed, based on previous work, that publicly accessible articles would be more highly accessed and thus more highly cited[35, 40, 41]. We did, however, find a positive association between citations and the interaction of public accessibility and JIF. This finding suggests that citation rates will increase when an article is publicly accessible *and* is published in a journal with a higher impact factor.

## Access counts

While faculty in academia value citation counts, citations take time to accrue[7]. Access counts, on the other hand, are more or less immediate and thus can serve as early indicators of post-publication impact[42]. Of the studied altmetrics outlets, we did not identify any relationship between altmetrics activity and access counts. However, we did identify that an HPE article’s public accessibility was positively associated with access counts.

From a practical standpoint, our finding on the positive relationship between public accessibility and access counts has potential implications for HPE researchers who are compelled, and are often incentivized, to disseminate their work as broadly as possible. Although we have no way of knowing if this relationship is causal, it does have potential implications for HPE researchers eager to increase readership. For example, *BMC Medical Education* is an open access journal, which means that articles are publicly available from publication into perpetuity. In contrast, *Academic Medicine* makes its articles publicly accessible on their website as soon as they are published ahead of print and then, once in print, provides “delayed access”[43], which enables public access to the full-text after a one-year embargo. Considering these differences and the findings observed here, it may behoove authors to reflect on a journal’s access policies when choosing the best home for a given article. In addition, this positive relationship could have implications for journals and their publishers, as they consider their own open access policies.

Unlike previous HPE research, as well as work from other fields [44], we did not observe a relationship between access counts and altmetrics (22). We were surprised by this finding and believe this indicates a need for additional research in this domain. In regards to Twitter and Facebook, we propose one possible contributing factor is that these tools do not connect with library subscription systems. Therefore, if a potential reader encounters a tweet to an article, they may choose not to access that article assuming they will either not have access or be asked to pay a fee. Article links leading to paywalls have been associated with user frustration and can condition some users to avoid clicking such links (under the expectation that such links lead to a request to pay)[45].

This study has several important limitations. First and foremost, due to the correlational nature of the study design, we have no way of knowing if the observed relationships are causal. Second, in our sample, we included all articles that were publicly accessible. In some cases, an article was publicly accessible because its author paid a processing charge or because the journal’s editor specifically selected it. In other situations, an article was publically accessible due to a journal policy (e.g., *Academic Medicine’s* policy). These differences make the associations between public accessibility and our outcomes more difficult to interpret. In addition, because we analyzed articles from a subset of HPE journals, our findings may have limited generalizability within the whole of HPE publishing. Thus, future studies should consider an even broader selection of HPE journals, as well as a more granular approach to determining potential differences related to an article’s specific type of public accessibility. Such data would be valuable to an HPE researcher attempting to make an educated decision on which journal best fits their needs.

Another limitation relates to access counts. In particular, we note that users often access full-text articles through means other than the publisher’s website or a university’s library. For example, SciHub, a controversial article service, is one of the top sources of full-text articles, with 6 million downloads in February 2017 alone [46]. Access to articles from this illegal, but incredibly popular, gateway were not captured in our study. As services like SciHub become a part of the academic landscape, researchers need to consider the effects of such literature sources.

## Conclusion

Researchers have a responsibility to disseminate their work broadly. Traditionally, dissemination in academia has been measured through citation counts and JIFs, two measures that have been criticized for a variety of notable reasons. In this study, we examined the relationships between altmetrics, citations, and access counts (among several other variables) and found that a number of altmetrics outlets are positively associated with citations. Furthermore, we found that public accessibility is positively related to article access. Given the scientific community’s evolving focus on dissemination, our findings have implications for a variety of HPE stakeholders, providing important insights into the factors that may improve an article’s citations and access.

